# Optimal sparsity in autoencoder memory models of the hippocampus

**DOI:** 10.1101/2025.01.06.631574

**Authors:** Abhishek Shah, René Hen, Attila Losonczy, Stefano Fusi

## Abstract

Storing complex correlated memories is significantly more efficient when memories are recoded to obtain compressed representations. Previous work has shown that compression can be implemented in a simple neural circuit, which can be described as a sparse autoencoder. The activity of the encoding units in these models recapitulates the activity of hippocampal neurons recorded in multiple experiments. However, these investigations have assumed that the level of sparsity is fixed and that inputs have the same statistics and, hence, that they are uniformly compressible. In contrast, biological agents encounter environments with vastly different memory demands and compressibility. Here, we investigate whether the compressibility of inputs determines optimal sparsity in sparse autoencoders. We find 1) that as the compressibility of inputs increases, the optimal coding level decreases, 2) that the desired coding level diverges from the observed coding level as a function of both memory demand and input compressibility, and 3) that optimal memory capacity is achieved when sparsity is weakly enforced. In addition, we characterize how sparsity and the strength of sparsity enforcement jointly control optimal performance. These results provide predictions for how sparsity in the hippocampus should change in response to environmental statistics and theoretical grounds for why sparsity is dynamically tuned in the brain.

## Introduction

The hippocampus clearly plays a fundamental role in memory consolidation. Several memory models of the hippocampus are based on the idea that efficiently compressing memories before they are stored can lead to a significantly higher memory capacity. Typically, theoretical studies on the memory capacity of artificial neural networks made a critical assumption about the nature of the memories that are stored: the patterns that represent memories were assumed to be random and uncorrelated (see, e.g. Hopfield (1982); Amit (1989); Benna and Fusi (2016); Fusi (2024)), mostly for convenience. However, most of our sensory experiences are highly correlated. To efficiently store these correlated experiences, it is essential to preprocess or recode the memory neural representations as suggested by previous research (Marr, 1971; Gluck and Myers, 1993; Treves and Rolls, 1994; McClelland and Goddard, 1996; Hasselmo and Wyble, 1997). Ideally, one would aim to extract the uncorrelated (and therefore incompressible) components of the new input patterns and store only those. This method of preprocessing and compression has been explicitly modeled in Benna and Fusi (2021), where the authors demonstrated 1) a significant improvement in memory performance and 2) that the neurons in the hippocampal model exhibit response properties that are similar to those observed in numerous experiments, including those of the place cells. Similar approaches have been recently studied in Recanatesi et al. (2018); Santos-Pata et al. (2021); Levenstein et al. (2024); Amil et al. (2024) in which the authors modeled the hippocampus either as a sparse autoencoder, a predictive network that is trained to predict the next neural state, or both.

In Benna and Fusi (2021), compression is achieved by modeling the hippocampus as a sparse autoencoder (Olshausen and Field, 1996, 1997), which is trained to reconstruct the memory to be recalled. Modeling the hippocampus as an autoencoder was originally proposed in Gluck and Myers (1993) and then studied in several other theoretical works (Treves and Rolls, 1994; McClelland and Goddard, 1996; Hasselmo and Wyble, 1997; Schapiro et al., 2017; Lian and Burkitt, 2020). In most of these models the representations are sparsified and orthogonalized in the intermediate layer of the autoencoder, making the memories more separable, which in turn facilitates their storage and reconstruction. Indeed, it is well known the memory capacity for sparse random patterns is significantly higher than in the case of dense patterns (Tsodyks and Feigel’man, 1988; Treves and Rolls, 1991). Moreover, orthogonalization is compatible with many experimental observations (see e.g. Sun et al. (2023); Boyle et al. (2024).

Here, we systematically studied how the level of sparseness imposed on a hippocampal autoencoder model determines memory capacity. We found that there is an optimal sparseness that depends on the total number of memories that have to be stored and, importantly, on their degree of compressibility.

## Results

### Model setup

Our memory benchmark is the same introduced in Benna and Fusi (2021): the set of patterns to be stored as memories is organized as an ultrametric tree (see e.g. Rammal et al. (1986) and Figure 1A). To generate correlated patterns, one starts with *A* uncorrelated random patterns, referred to as the ancestors at the top of the tree. In these patterns, each neuron is either active or inactive with equal probability, similar to the Hopfield model (Hopfield, 1982). From each ancestor, one can generate *K* descendants by randomly resampling the activation states of the neurons in the ancestors with a probability *γ. K* is known as the branching ratio of the tree. The total number of descendants will be *P* = *A* × *K*, and these are the patterns intended for storage in memory. Patterns that descend from the same ancestor are correlated, while those generated from distinct ancestors are uncorrelated. This basic method for generating correlations between patterns has been studied to extend the Hopfield model to correlated attractors (Feigelman and Ioffe, 1996; Gutfreund, 1988; Fusi, 1995).

**Figure 1.**
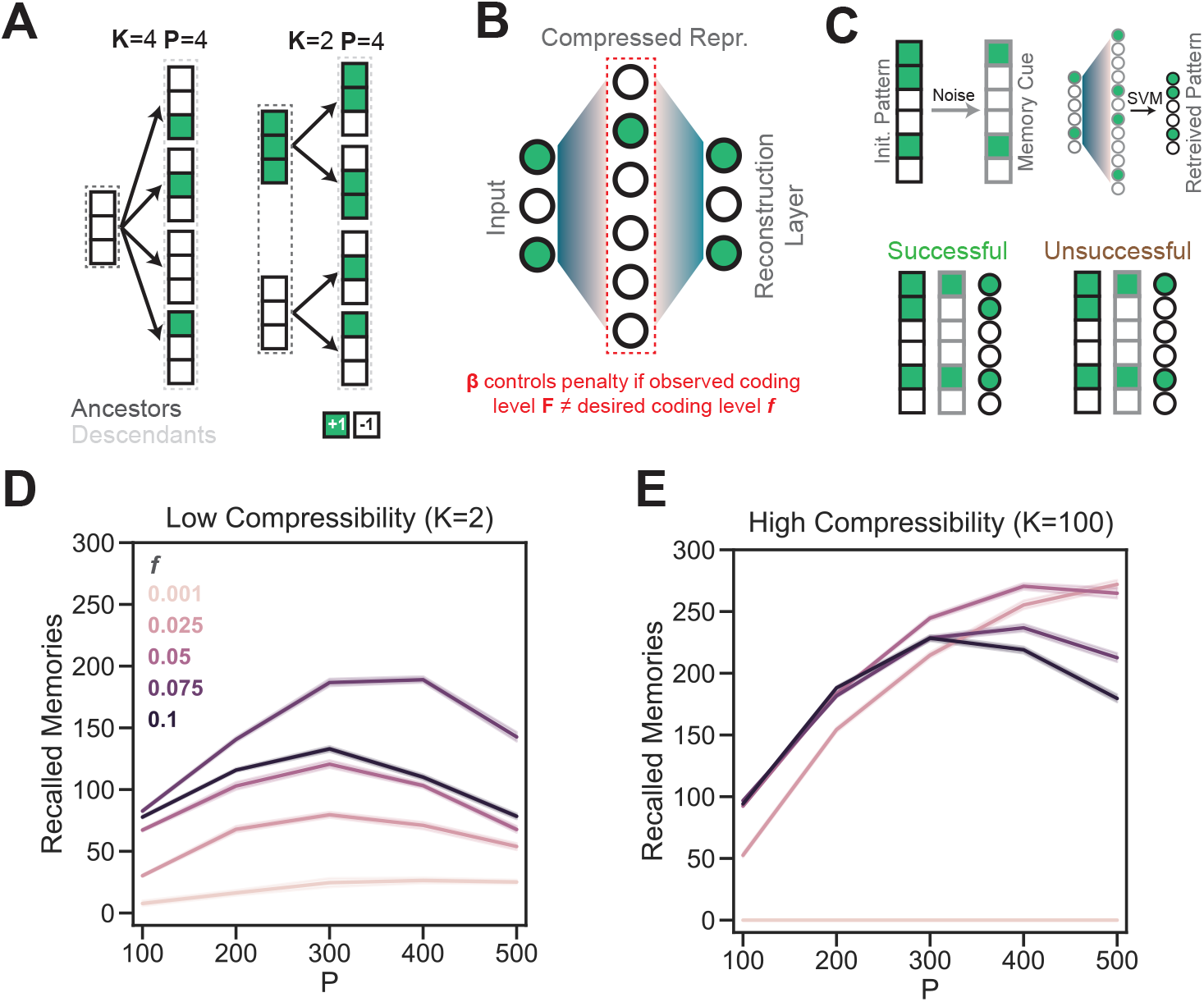
Sparse autoencoders can be trained with varying desired coding levels and input compressibilities. (**A**) Ultrametric trees are used to generate compressible inputs for autoencoders. On the left, a highly compressible environment is illustrated, where a total of *P* = 4 patterns are generated by randomly noising a single ancestor with a branching ratio *K* = 4. On the right, a less compressible environment is generated by using a lower branching ratio, such that to generate four patterns, two random ancestors are needed. (**B**) The sparse autoencoder takes in a single pattern and is trained to reconstruct that pattern, with the constraint that encoding units should have a certain coding level *f*. If the observed coding level *F* does not line up with the desired coding level, a penalty is applied during training, the strength of which is controlled by the parameter *β* (set to 1 in all plots in this figure). (**C**) To test whether networks accurately store memories, the original patterns that the network was trained on are noised. These noised inputs are then presented to the network, and a support vector machine (SVM) is used to decode the original pattern. If the decoded pattern has high overlap with the original pattern, it is considered a successful recollection. If the decoded pattern does not have high overlap - for instance, if it simply recreates the noised pattern - the pattern is considered to be unsuccessfully retrieved. (**D**) Memory performance at low compressibility, across various numbers of patterns *P* in the training set provided to the autoencoder. Hues differentiate models trained with different desired coding levels. (**E**) Same as D, but with high input compressibility.

For a fixed set of *P* total patterns, increasing the number of ancestors (and therefore decreasing the branching ratio) used to generate the final patterns leads to a decrease in the compressibility of the final set of patterns. For simplicity in the rest of the paper, we solely refer to the branching ratio *K* instead of the number of ancestors, as an increase in *K* corresponds to an increase in the compressibility of a set of patterns. Only descendants were fed as inputs to networks.

To impose sparsity, a penalty was added to the loss function if the average activation in the hidden layer deviated from the desired coding level *f*. The relative contribution of this component to the loss was controlled by the parameter *β* (Fig. 1B). To test for memory performance, it was necessary to determine how well networks could recall learned patterns from degraded inputs (Fig. 1C). To do this, memory cues were generated for the training set by noising each each training pattern. A support vector machine (SVM) was then trained to decode training patterns from hidden layer encodings. This trained SVM was then tested on the encodings of the noised patterns. Memory performance was defined as the proportion of patterns that were successfully reconstructed by the SVM. As has been previously described (Benna and Fusi, 2021), varying the desired coding level *f* led to shifts in the number of recalled memories (Fig.1D) when patterns had low compressibility, with the greatest number of recalled patterns at an *f* of 0.075. Interestingly, when patterns had high compressibility *K*, the greatest number of recalled memories resulted from networks trained with a desired coding level of 0.05 instead of 0.075, indicating that the optimal *f* for memory performance is contingent on the compressibility of patterns to be memorized (Figure1E).

### Optimal sparsity is tuned to the compressibility of inputs

To rigorously understand the dependence of optimal sparsity on the compressibility of inputs, multiple networks were trained with the number of patterns fixed at *P* = 400. In networks with strong enforcement (*β* = 5) of sparsity, an increase in the compressibility of inputs led to a decrease in the desired coding level (i.e., an increase in the sparsity) that yielded maximum memory performance (Fig. 2A). Since memory performance was tested on memory cues that were noised versions of the original inputs, we were interested in whether the coding level of noised memory cues was aligned with the desired coding level with which networks were trained. We were also interested in whether the compressibility-dependent shift in the optimal desired coding level would persist for the observed coding level of memory cues. While there was an increase in the optimal observed coding level when the branching ratio *K* decreased from 100 to 25, there was an unexpected decrease in the optimal observed coding level when the branching ratio further decreased to *K* = 2 (Fig. 2B).

**Figure 2.**
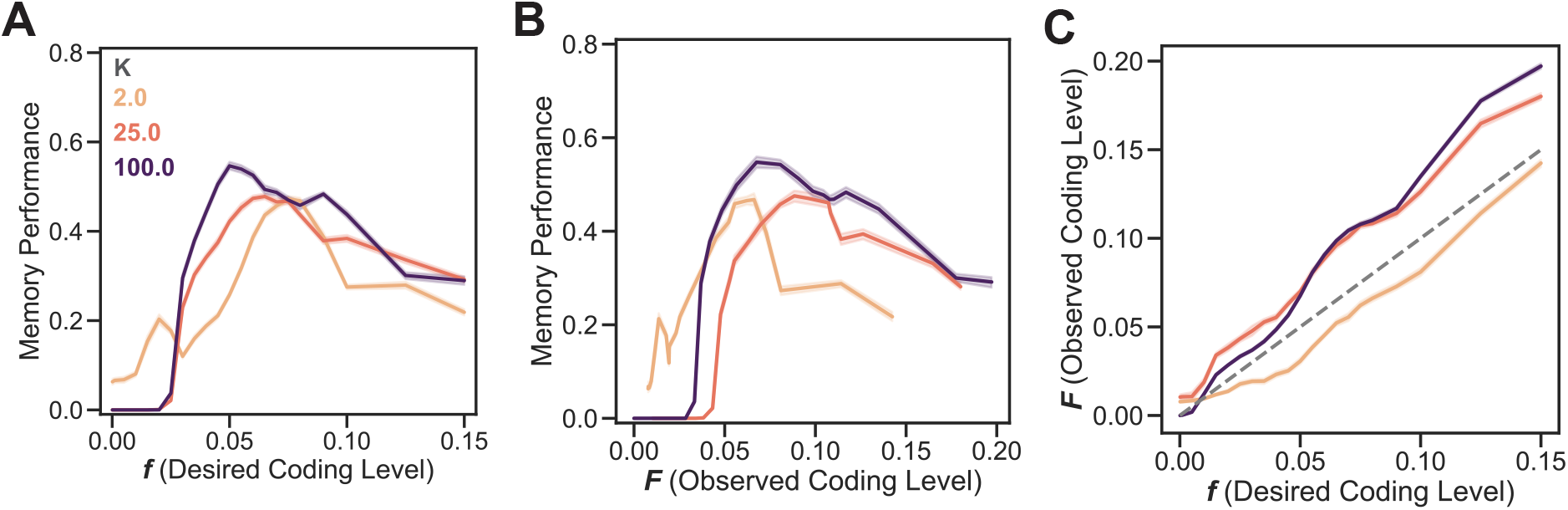
Networks with strong (*β* =5) enforcement of sparsity have optimal desired coding levels that vary as a function of input compressibility. (**A**) Memory performance as a proportion of successfully retrieved memories for a fixed number of inputs *P* = 400, across various compressibilities, as a function of the desired coding level. (**B**) Similar to A, but instead of the desired coding level, the average observed coding level *F* for the noised memory cues for each network is plotted. **C** The observed coding level for each network as a function of the desired coding level provided during training. The dashed line is the identity line.

As these results implied, the observed coding level for encodings of noised memory cues was roughly linear for *K* = 25, 100 (Fig. 2C). However, for *K* = 2, the observed coding level was consistently lower than the desired coding level (Fig. 2C). These results indicate that the optimal desired coding level for sparse autoencoders with strong sparsity enforcement has an inverse relation to the compressibility of inputs. Additionally, the coding level of noised memory cues does track the desired coding level, but the strength of this relation is contingent on the compressibility of inputs. Given that the coding levels of the noised cues strayed from the line of identity, we were interested in whether weaker enforcement of sparsity would allow networks to tend towards a more optimal observed sparsity level.

### Networks maintain memory performance at higher coding levels with weaker sparsity enforcement

Similar to as in Figure 2, models were trained across a range of *K* and *f*, but this time with weak sparsity enforcement (*β* = 0.25). While a clear optimal desired coding level was present for *K* = 100, for lower compressibilities memory performance plateaued from the lowest tested desired coding levels (*f* = 0.001), all the way up to *f* = 0.075 (Fig.3A). When considering the observed coding level, it became clear that networks tended towards coding levels that were far greater than desired coding levels (Fig.3B,C). In spite of this, networks tended towards a lower optimal observed coding level for inputs with higher compressibility, mimicking the trend seen in networks with stronger enforcement of sparsity. However, this effect was far slighter than in models with strong sparsity enforcement, and a return to lower optimal coding levels for low compressibilities was not seen as was the case in Fig. 2B. These results indicate that weaker enforcement of sparsity induces networks to tend towards the same optimal coding level for multiple desired coding levels, leading to a ‘plateau’ in performance and observed coding levels across multiple desired coding levels.

**Figure 3.**
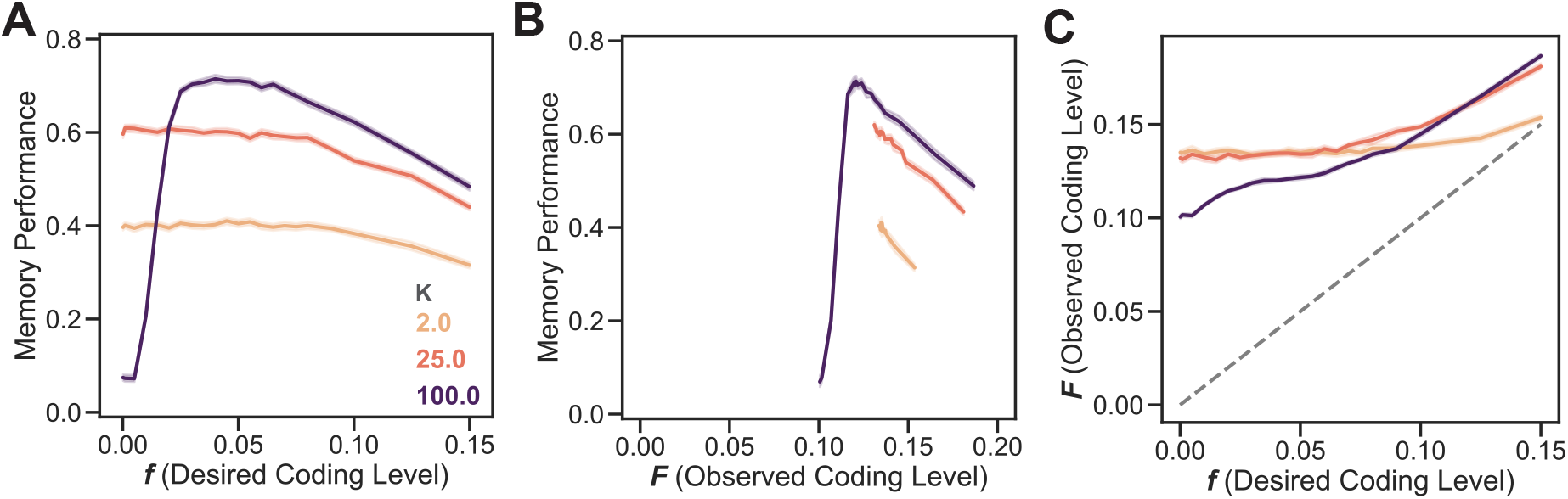
Networks with weaker (*β* =0.25) enforcement of sparsity develop distinct coding levels after training as a function of input compressibility. (**A**) Similar to Fig. 2A, but with weaker enforcement of sparsity. (**B**) Similar to A, but instead of the desired coding level, the average observed coding level *F* for memory cue encodings for each network is plotted. Note that lines are not censored - no observed coding levels fall below certain thresholds for each set of networks. **C** The observed coding level for each network as a function of the desired coding level provided during training. The dashed line is the identity line.

### Weak enforcement of sparsity generally increases memory performance

Paradoxically, while maintaining sparsity appears to be important for boosting memory capacity (Fig.1D), weaker enforcement of sparsity did not degrade memory performance (Benna and Fusi, 2021). In fact, strong enforcement of sparsity led to degraded memory performance across most tested coding values and input compressibilities (Fig.4A,B). However, depending on the total number of memories in the inputs, the optimal enforcement of sparsity also varied. At low memory capacities (*P* = 100), performance was generally uniform across all sparsity enforcement strengths at *f* > 0.05, with a slight advantage for strong sparsity enforcement when compressibility was also high (Fig.5A,C). However, with higher memory demand, moderate sparsity enforcement was optimal at low compressibility, but both weak and moderate sparsity enforcement were optimal at higher compressibilities (Fig.5B,D). These data indicate that optimal sparsity enforcement is tuned not only to the compressibility of inputs but also to the total number of memories that a network must store.

**Figure 4.**
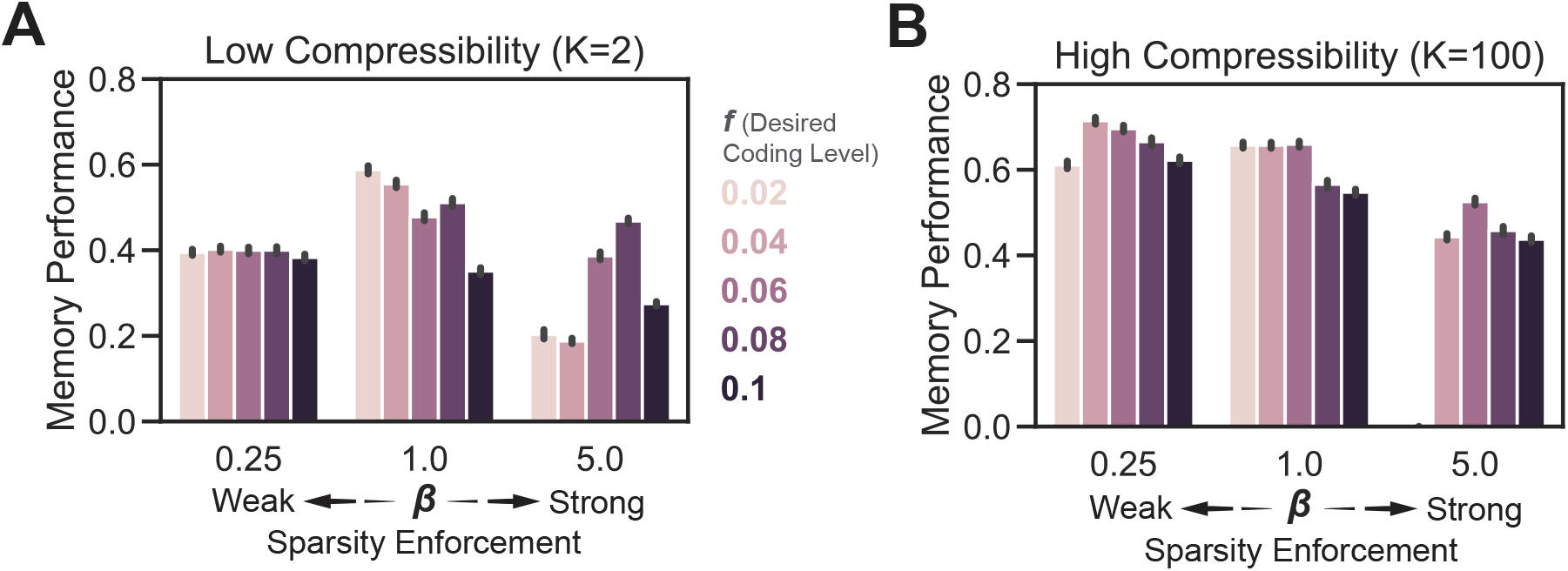
Weaker enforcement of sparsity is optimal for memory performance across all tested desired coding levels. (**A**) Memory performance is plotted for a given desired coding level, split by *β*, for a weakly compressible set of patterns. (**B**) Same as (A), but with a highly compressible environment. *P* = 400 was used for all networks plotted here. Bars are 95% CI.

**Figure 5.**
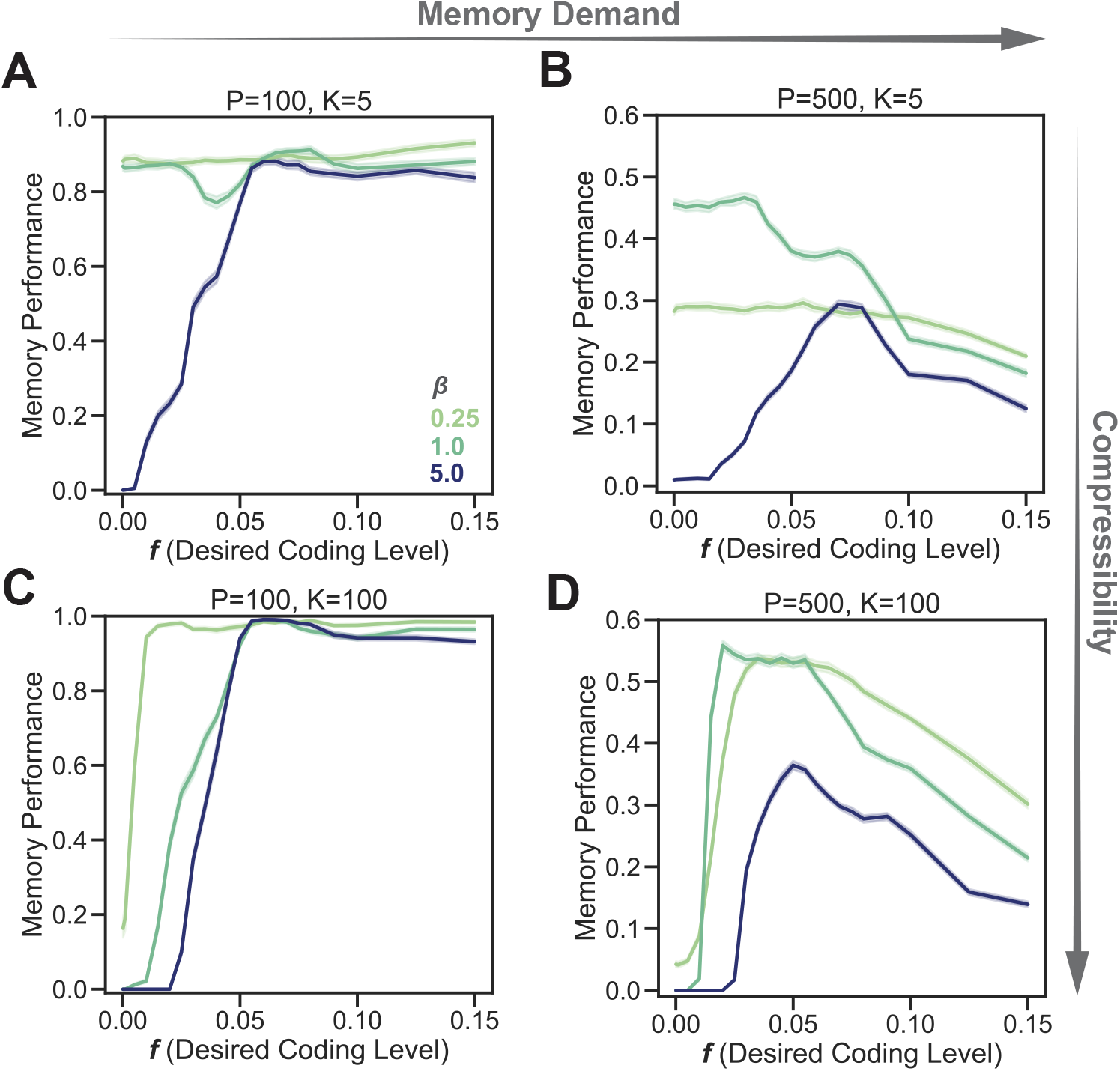
Optimal sparsity enforcement and sparsity are also contingent on memory demand. (**A**) Memory performance as a proportion of memories in the input set for low memory demand and low compressibility as a function of the desired coding level. Hues represent networks trained with different *β*. (**B**) Same as (A), but for higher memory demand. (**C-D**) Same as (A) and (B), but with high input compressibility.

## Discussion

### Optimal sparsity is contingent on compressibility and memory demand

Across all models and parameters tested, there were clear optimal coding levels where memory performance was maximal. Optimal coding levels were present both when considering the desired coding level and the observed coding level during the memory task (Fig.2A,B,3A,B). The optimal desired coding level was contingent on the compressibility of the set of training patterns, with a decrease in the optimal coding level as compressibility increased. While strong enforcement of sparsity was able to control the coding level during memory tests, it was also possible to weakly enforce sparsity to observe what coding levels the network would evolve towards during training. Interestingly, weaker enforcement of sparsity led to higher memory performance, which was accompanied by an increase in observed coding levels (Fig.4A,B,3C). These results indicate that while maintaining sparsity is important for memory performance, penalizing networks too harshly for maintaining denser encodings prevents the training of an encoder that can properly generalize to noisy inputs. It is important to note here that simply achieving a higher coding level does not lead to better performance on the memory task: when sparsity was strongly enforced, networks trained with higher desired coding levels showed performance that was half of peak performance across all models (Fig.2B,C).

Memory demand was also important for determining the strength of sparsity enforcement: when networks were trained on fewer patterns, they generally achieved near-perfect memory performance across all tested sparsity enforcement strengths, but when memory demand increased, it became important to maintain weaker sparsity enforcement. (Fig.5A-D). As a whole, these results suggest that sparsity is tuned to the compressibility of inputs. However, it is also important for networks to be given leeway to tend towards a a higher coding level to maximize performance on the memory task, where it is important that encodings are robust to noise.

### Enforcing sparsity with alternative measures

In order to maintain a differentiable loss function, we utilized a loss function that penalized deviations in the sparsity of an encoding vector from the desired sparsity *f* by simply taking the average across activations in the encoding layer (Eq.1). There are other ways of enforcing sparsity: for instance, Kullback-Leibler divergence and similar measures of the distribution of activations are often used in sparse autoencoders, but they generally measure sparsity per-neuron (e.g., the coding level measures the proportion of patterns in the entire set that a given neuron is active for), rather than per-encoding (e.g. the proportion of active units in the encoding for a single pattern) (Nair and Hinton, 2009). Alternative approaches that can be applied per-vector include forcing sparsity by choosing the k-strongest activations, or *L*^0^ norm minimization, which explicitly measures sparsity but requires approximation to be used with gradient descent (Makhzani and Frey, 2014; Willmore and Tolhurst, 2001). It will be important for future work to test whether the dependency of optimal coding level on input compressibility is consistent across these alternate sparsity measures. We also utilized *L*^2^-approximate weight regularization in our models to prevent trivial readout of sparse codes. It is possible that this could play an important role in training an encoder that generalizes, and that the interplay between the sparsity loss and the impact of regularization has an important role in shaping these networks (Steck, 2020).

### Flexible sparsity in the brain

Our model allowed for both a desired sparsity (*f*) and the relative strength of sparsity enforcement (*β*) to be strictly controlled. How might the brain dynamically tune the level of sparsity in order to optimally match input statistics? In the hippocampus, a sparse code is maintained in the dentate gyrus (DG) via strong lateral inhibition (Espinoza et al., 2018). In addition to this lateral inhibition, a diverse set of inhibitory interneurons appear to be tuned to behavior, novelty, and stress (Nitz and McNaughton, 2004; Schoenfeld et al., 2013). Lateral inhibition has already been shown to play an important role in biologically plausible implementations of autoencoders (Pehlevan and Chklovskii, 2015; Pehle-van et al., 2018; Sengupta et al., 2018; Benna and Fusi, 2021). Both lateral and feedforward inhibition might be tuned dynamically to rapidly optimize DG coding level in response to acute input compressibility. Recent evidence also suggests that sparsity can be driven at longer timescales by neurogenesis in the DG, which in turn is tuned to the richness (and perhaps compressibility) of environments, as well as to stress (Luna et al., 2019; McHugh et al., 2022). Therefore, neurogenesis might provide a slower mechanism to tune sparsity as an animal integrates over many experiences.

### Experimental predictions

Our results indicate that in more compressible environments, the DG would favor a sparser code. While measuring the compressibility of an animal’s experience can be difficult, it is known that when animals are stressed or in an unenriched environment, the sparsity of the DG decreases (Karst and Joëls, 2003). Traditionally, it is thought that this favors pattern completion rather than pattern separation, allowing for generalization of experience (Treves et al., 2008). While environments with low richness might appear to be highly compressible, in our framework, they might be better understood as environments with low memory demand (Fig.5A-D). In these environments, it is likely that higher coding levels do not penalize memory performance, as our model predicts. In contrast, in richer environments, memory demand is likely higher, which drives down the optimal coding level. To test the effects of environmental compressibility on the coding level of the DG, it will be necessary to strictly control the statistics of experience, likely via virtual reality environments, while keeping the total memory demand fixed.

As a whole, our findings indicate that sparsity in the DG is likely tuned to both the compressibility and total memory demand of inputs and that changes in DG coding level in response to behavior, cognitive state, and environmental statistics could be adaptive responses to optimize memory performance.

## Methods

### Generating compressible pattern sets

Ultrametric trees were used to generate compressible sets of patterns as described in Benna and Fusi (2021). To generate *P* patterns with a branching ratio of *K, A* = *P/K* total 300-unit ancestors were generated by randomly sampling each unit from the Rademacher distribution, such that each unit had an equal chance of being -1 or 1. *K* descendants were then generated from each ancestor pattern by resampling each unit w/probability *γ* = 0.4.

### Training sparse autoencoders

All networks were trained in Python (3.11.8) using Pytorch (2.1.1). Networks consisted of a fully connected input (300 units), encoding (600 units), and output (300 units) layer, with sigmoid activations between the input and encoding layers. To maintain sparsity, the average activation value in the encoding layer was used as a proxy for the coding level, and a penalty was applied if this value differed from the desired sparsity *f*, weighted by the coefficient *β*. All networks were trained to reconstruct their input units by utilizing mean-squared error (MSE) loss, such that the final joint loss function for a batch of patterns *X* was: (Equation 1).

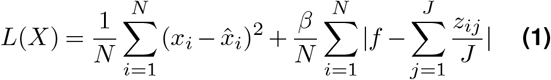

Where *N* is the number of patterns in the batch, *x*_*i*_ is the *i*-th pattern in *X*, 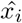 is the output from the network for an input *x*_*i*_, *β* is a constant, *f* is the desired coding level as a proportion, *J* is the number of units in the encoding layer and *z*_*ij*_ is the activation of the *j*-th unit in the encoding for an input *x*_*i*_. Adam was used as the optimizer for all modeling, with weight regularization performed by setting weight_decay to 0.00001. The initial learning rate was set to 0.001, and training was performed for 3000 epochs or until the total loss dropped beneath 0.01.

### Testing memory performance

Refer to Figure 2C for a graphical description of the memory performance procedure. First, memory cues were generated by inverting the value of each unit in the training set of patterns with probability 0.2, such that, on average, memory cues were 80% similar to patterns in the training set. Each memory cue was then presented to the trained network, and the hidden layer encoding was generated for each cue. A set of SVMs (Scipy 1.11.4, LinearSVC, C=0.005) was then trained to decode input patterns from the binarized hidden layer encodings of the original training patterns. Each SVM was trained to decode a single input unit from the entire encoding vector, and binarization was performed by mapping any value in the encoding layer greater than 0.5 to 1, and less than 0.5 to -1. These trained SVMs were then tested on the binarized encodings from the memory cues. A pattern was considered correctly recalled if more than 90% of units in the decoded pattern overlapped with the original pattern from the training set. Observed coding levels for the memory cues were obtained by binarizing the encodings using the same method mentioned above. Reported memory performance corresponds to the proportion of correctly recalled patterns from the full set of noised memory cues.

## Acknowledgements

We thank Jack Lindsay, James Priestley and Lorenzo Posani for many useful discussions. A.S. and S.F. were supported by the Gatsby Charitable Foundation (GAT3708), the Kavli Foundation and the Swartz Foundation. A.S., S.F., R.H., and A.L. were supported by the National Institute on Aging (1RF1AG080818). A.S. was also supported by the National Science Foundation Graduate Research Fellowship (DGE-2036197).

